# Identification of novel mutations in LprG *(rv1411c), rv0521, rv3630, rv0010c, ppsC, cyp128* associated with pyrazinoic acid/pyrazinamide resistance in *Mycobacterium tuberculosis*

**DOI:** 10.1101/249201

**Authors:** Wanliang Shi, Jiazhen Chen, Shuo Zhang, Wenhong Zhang, Ying Zhang

**Affiliations:** Department of Molecular Microbiology and Immunology, Bloomberg School of Public Health, Johns Hopkins University, Baltimore, MD 21205, USA; Department of Infectious Diseases, Huashan Hospital, Fudan University, Shanghai 200040, China

## Abstract

There is currently considerable interest in understanding the mechanisms of action of pyrazinamide (PZA), a critical frontline tuberculosis (TB) drug that plays a unique role in shortening TB therapy due to its unique activity against *Mycobacterium tuberculosis* persisters that are not killed by other TB drugs.^1,2^ Despite the importance of PZA in the treatment of both drug susceptible and drug-resistant TB and its simple structure, its mechanisms of action are complex and are not well understood.^1,2^ PZA is a prodrug that is converted to the active form pyrazinoic acid (POA) by nicotinamidase/pyrazinamidase (PZase) encoded by the *pncA* gene,^3^ whose mutation is the most common mechanism of PZA resistance in *M. tuberculosis.*^*3-5*^ However, some low level PZA-resistant strains (MIC=200-300 μg/ml, pH6.0) do not have mutations in the *pncA* gene.^5,6^ Recent studies have identified *rpsA*, which encodes the ribosomal protein S1 involved in both translation and trans-translation process, as a target of PZA,^7^ where its mutations are associated with PZA resistance from clinical isolates. In addition, mutations in *panD* encoding aspartate decarboxylase were identified as a new mechanism of PZA resistance from in vitro mutants resistant to PZA and the PanD protein was found to be another target of PZA.^8,9^ *panD* mutations were initially found in mutants resistant to PZA ^8^ and then in mutants resistant to POA. ^9,10^ It is worth noting that previously POA-resistant mutants could not be isolated at acid pH which is required for higher activity of PZA against *M. tuberculosis.* However, we were able to successfully isolate POA-resistant mutants with high POA concentrations at close to neutral pH (pH 6.8),^9^ which led to discovery of new genes involved in POA and PZA resistance. For example, *clpC1*, which was also isolated from mutants resistant to PZA,^11^ was identified in mutants resistant to POA.^12^

---

However, in our previous study, only a small number of 30 POA-resistant mutants of *M. tuberculosis* were analyzed, where all the mutations were mapped to the *panD* gene.^9^ Since some PZA-resistant strains such as strain 9739 ^5^ do not have any mutations in the known genes *pncA*, *rpsA*, *panD*, or *clpCl* involved in PZA resistance (Shi W and Zhang Y, unpublished), this would indicate that new mechanisms of POA/PZA resistance still exist. In this study, to identify possible new mechanisms of PZA resistance, we characterized an additional 109 POA-resistant mutants (MIC=200 μg/ml POA, pH6.8) of *M. tuberculosis* H37Ra (MIC<100 μg/ml POA) derived from our previous study.^9^ Briefly, the genomic DNA from the 109 POA-resistant mutants was extracted and subjected to PCR DNA sequencing of the known genes *pncA*, *rpsA*, *panD*, *clpCl* involved in PZA resistance. It is noteworthy that 101/109 (93%) of POA-resistant mutants had *panD* mutations, with 75 of the *panD* mutants having the same dominant M117I mutation as previsouly identified.^9^ However, 8 POA-resistant mutants did not have *panD* mutations, and 4 of the mutants had the same mutation in *clpC1* where A271G nucleotide change caused amino acid substitution S91G. It is noteworthy that the *clpC1* mutation S91G is different from our previsouly identified mutation (G99D)^11^ and those by Gopal *et al‥*^12^ However, the remaining 4 POA-resistant mutants were also resistant to PZA, which are 2-4 fold more resistant (50-100 μg/ml PZA, pH5.8) than parent strain *M. tuberculosis* H37Ra (25 μg/ml PZA, pH5.8); but these 4 POA-resistant mutants did not have any mutations in the known PZA resistance genes *pncA*, *rpsA*, *panD*, or *clpC1.* Whole genome sequencing of the 4 POA-resistant mutants was performed as we described previously,^8^ and sequence analysis revealed that they all had the same two mutations *rv1411c* (Lipoprotein LprG C672T change leading to W224STOP) and *rv0521* (Methyltransferase, C295T causing amino acid substitution R99W) (Table 1). Sanger sequencing of the PCR products confirmed the mutations identified by the whole genome sequencing. However, Strain 3G1 had only the two identical mutations in LprG *(rv1411c)* and *rv0521* while the remaining 3 mutants had additional low percentage of heterogeneous mutations in *rv3630* (hypothetical protein), *rv0010c* (hypothetical protein), *ppsC* (phenolpthiocerol synthesis type-I polyketide synthase C) and *cyp128* (cytochrome P450 128) (Table 1). Strain 3H4 had a point mutation T199C, causing amino acid change T67A in cytochrome P450 128 (*cyp128*, Rv2268c), which encodes a heme-thiolate monooxygenase.

**Table 1.**
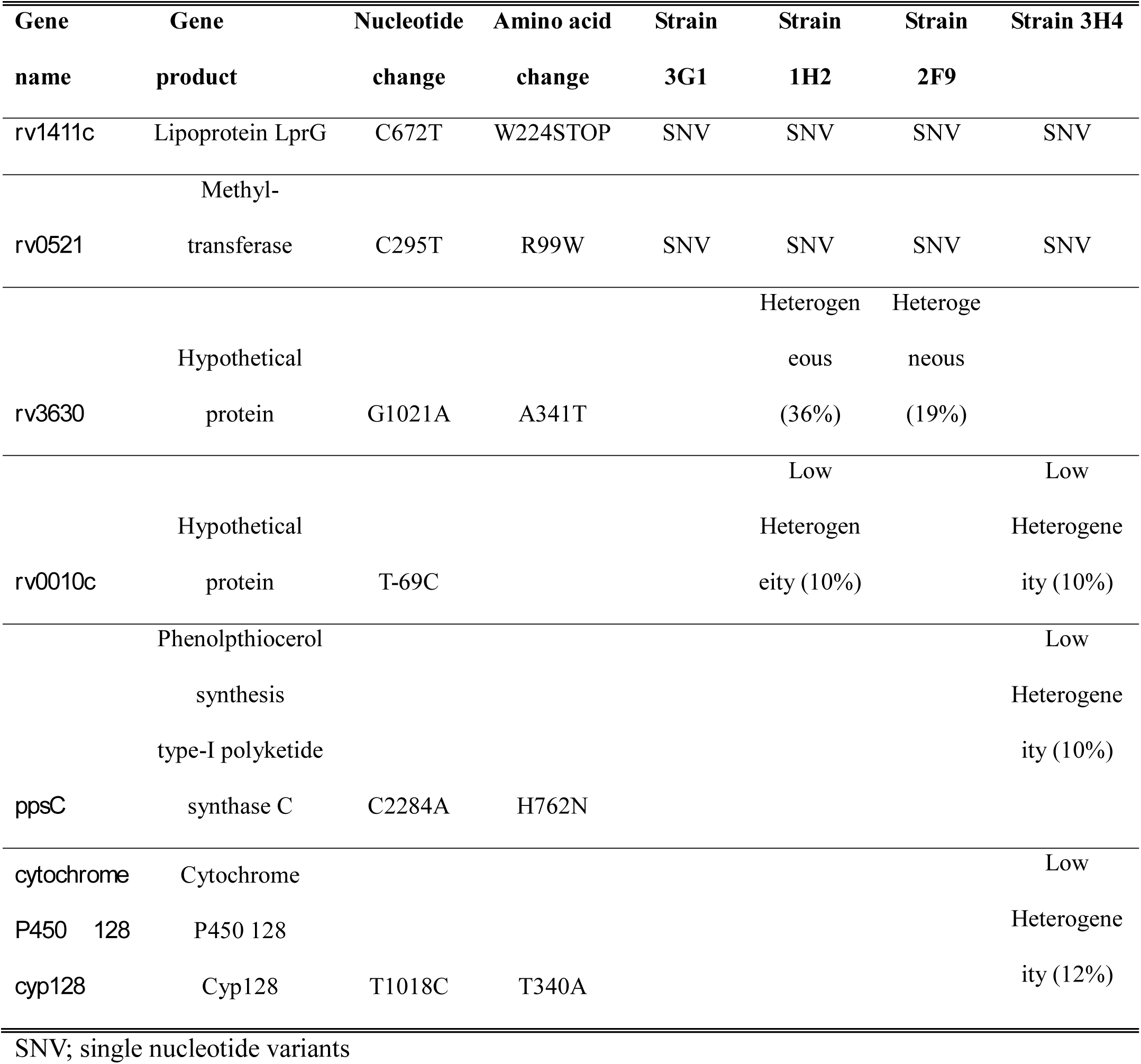
Mutations identified in the POA-resistant mutants of *M. tuberculosis* that do not have known PZA-resistance mutations

LprG-Rv1410 has been shown to participate in triacylglyceride (TAG) transport in *M. tuberculosis* as loss of its function leads to intracellular TAG accumulation, and is critical for regulating bacterial growth and metabolism during carbon starvation and infection in mice.^13^ The finding that all 4 POA-resistant mutants have the same W224Stop mutation in LprG suggests that the non-functional LprG will lead to higher TAG accumulation and thus causing higher metabolism, which is known to be a condition that antagonizes PZA/POA activity.^2^ This provides a plausible explanation for loss of function mutation in LprG being a likely cause for PZA/POA resistance. However, the role of the mutation in *rv0521* which encodes a putative methyltransferase in causing POA resistance is less clear, and it is not known why the 4 POA-resistant mutants all had the same 2 mutations in LprG and Rv0521. Since one POA-resistant mutant 3G1 has only the two identical mutations and no other mutations, it can be inferred that either LprG or Rv0521or both must be involved in POA/PZA resistance. Future studies are needed to address the contribution of each individual gene *lprG* and *rv0521* in causing POA/PZA resistance and their possible role in the mode of action of PZA.

However, the contribution of the low abundance mutations in *rv3630* (hypothetical protein), *rv0010c* (hypothetical protein), *ppsC* (phenolpthiocerol synthesis type-I polyketide synthase C) and *cyp128* (cytochrome P450 128) (Table 1) is less clear. Previous studies have found mutations in *ppsA-E* locus involved in synthesis of virulence factor phthiocerol dimycocerosate (PDIM) are associated with PZA resistance,^10^ but the role of the *pps* operon gene mutations including *ppsC* mutations in PZA resistance remains to be determined. In *M. tuberculosis*, cytochrome P450 proteins have many functions such as virulence, persistence, cholesterol metabolism, lipid degradation, regulator, resistance to azole drugs.^14^ The *M. tuberculosis cyp128* is one of three essential cytochrome P450s (*cyp121*, *125*, *128*) in *M. tuberculosis*, and is part of an operon with *rv2269c*, and *sft3*, a sulfotransferase involved in biosynthesis of sulfomenaquinone (SMK), and disruption of SMK synthesis affects *M. tuberculosis* respiration and fitness.^15^ *cyp128* is up-regulated during starvation and is essential for survival in mice.^14^ Although we found that a single mutation in *cyp128* (T199C) causing amino acid change T67A associated with POA and PZA resistance, future studies are needed to confirm the role of cytochrome P450 protein Cyp128 in mechanisms of action and resistance of PZA. While a causative role of the low abundance gene mutations cannot be ruled out, it is likely that these mutations may serve to elevate resistance levels or allow the mutant to adapt to the two mutations in LprG and methyltransferase. These possibilities remain to be tested in future studies.

Despite the complexity of the mode of action of PZA, significant progress has been made in recent years in our understanding of this peculiar and yet critical drug, as PZA has been shown to interfere with multiple targets including trans-translation (RpsA), energy production (PanD), protein degradation (mediated by ClpC1), all of which are involved in persister survival in *M. tuberculosis.* This study identified novel genes associated with POA/PZA resistance, which may shed new light on the mechanisms of action of PZA. Future studies are needed to address the role of the identified mutations in PZA resistance and how they might be involved in the mode of action of PZA in M *tuberculosis*.

## ACKNOWLEDGEMENTS

This study was supported in part by NIH grant AI099512 and the National Science Foundation of China (81772231 and 81572046).

